# Ultra-deep sequencing differentiates patterns of skin clonal mutations associated with sun-exposure status and skin cancer burden

**DOI:** 10.1101/2020.01.10.902098

**Authors:** Lei Wei, Sean R. Christensen, Megan Fitzgerald, James Graham, Nicholas Hutson, Chi Zhang, Ziyun Huang, Qiang Hu, Fenglin Zhan, Jun Xie, Jianmin Zhang, Song Liu, Eva Remenyik, Emese Gellen, Oscar R. Colegio, Michael Bax, Jinhui Xu, Haifan Lin, Wendy J. Huss, Barbara A. Foster, Gyorgy Paragh

**Author notes:** These authors contributed equally (Lei Wei, Sean R. Christensen, Wendy J. Huss, Barbara A. Foster, Gyorgy Paragh). Email addresses: LW SC MF JG NH CZ ZH QH FZ JX JZ SL ER EG OC MB JX HL WJH BAF GP.

## Abstract

Non-melanoma skin cancer is the most common human malignancy and is primarily caused by exposure to ultraviolet (UV) radiation. The earliest detectable precursor of UV-mediated skin cancer is the growth of cell groups harboring clonal mutation (CM) in clinically normal appearing skin. Systematic evaluation of CMs is crucial to understand early photo-carcinogenesis. Previous studies confirmed the presence of CMs in sun-exposed skin. However, the relationship between UV-exposure and the accumulation of CMs, and the correlation of CMs with skin cancer risk remain poorly understood. To elucidate the exact molecular and clinical effects of long-term UV-exposure on skin, we performed targeted ultra-deep sequencing in 450 individual-matched sun-exposed (SE) and non-sun-exposed (NE) epidermal punch biopsies obtained from clinically normal skin from 13 donors. A total of 638 CMs were identified, including 298 UV-signature mutations (USMs). The numbers of USMs per sample were three times higher in the SE samples and were associated with significantly higher variant allele frequencies (VAFs), compared with the NE samples. We identified genomic regions in *TP53, NOTCH1* and *GRM3* where mutation burden was significantly associated with UV-exposure. Six mutations were almost exclusively present in SE epidermis and accounted for 42% of the overall difference between SE and NE mutation burden. We defined Cumulative Relative Clonal Area (CRCA), a single metric of UV-damage calculated by the overall relative percentage of the sampled skin area affected by CMs. The CRCA was dramatically elevated by a median of 11.2 fold in SE compared to NE samples. In an extended cohort of SE normal skin samples from patients with a high- or low-burden of cutaneous squamous cell carcinoma (cSCC), the SE samples in high-cSCC patients contained significantly more USMs than SE samples in low-cSCC patients, with the difference mostly conferred by mutations from low-frequency clones (defined by VAF≤1%) but not expanded clones (VAF>1%). Our studies of differential mutational features in normal skin between paired SE/NE body sites and high/low-cSCC patients provide novel insights into the carcinogenic effect of UV exposure, and indicate that CMs might be used to develop novel biomarkers for predicting cancer risk.

**Significance statement:** In UV radiation exposed skin, mutations fuel clonal cell growth. We established a sequencing-based method to objectively assess the mutational differences between sun-exposed (SE) and non-sun-exposed (NE) areas of normal human skin. Striking differences, in both the numbers of mutations and variant allele frequencies, were found between SE and NE areas. Furthermore, we identified specific genomic regions where mutation burden is significantly associated with UV-exposure status. These findings revealed previously unknown mutational patterns associated with UV-exposure, providing important insights into UV radiation’s early carcinogenic effects. Additionally, in an extended cohort, we identified preliminary association between normal skin mutation burden and cancer risk. These findings pave the road for future development of quantitative measurement of subclinical UV damage and skin cancer risk.

## Background

Ultraviolet (UV) light is responsible for over 5 million cases of skin cancer annually in the US, which is more human malignancies than all other environmental carcinogens combined^1,2^. In mammals, nucleotide excision repair eliminates UV-mediated DNA lesions, but this mechanism of repair is error prone resulting in frequent mutations^3^. The preferential location of UVB induced DNA lesions results in a specific pattern of so-called UV signature mutations at dipyridine sites (C>T, CC>TT)^4^. In most skin cancers, including cutaneous squamous cell carcinoma (cSCC), the burden of UV signature driver mutations is high^4,5^. While some cSCC arise from visible precancerous lesions known as actinic keratoses (AKs), many cSCC arise in apparently “normal” skin areas from precursors that are clinically invisible^6^. Therefore, clinically visible precursors are an ominous sign but not a sensitive early measure of photocarcinogenesis.

*TP53* mutations are among the most common driver mutations in cSCC, and are also detected by immunohistochemistry in aged normal skin^7,8^. These UV-induced *TP53* mutations facilitate clonal expansion of cells harboring them and therefore behave as early clonal mutations (CMs)^9^. For two decades *TP53* mutant keratinocyte cell clones were considered the earliest manifestations of skin carcinogenesis^7,8,10^. Because p53 clonal immunopositivity could not be efficiently quantified in human skin, detection of mutant *TP53* for assessment of photocarcinogenesis in clinical dermatology practice has been unattainable. The low relative abundance of clonal DNA previously limited efficient detection of early mutated cell groups. However, with improved high throughput sequencing technology we have finally reached the lower end of this threshold and efficient detection of rare mutations in normal tissue is becoming feasible in recent studies by others and us using deep bulk sequencing or single cell DNA sequencing ^11-16^. In exploratory analyses, CMs were found to be abundant in clinically normal skin from sun-exposed sites in *NOTCH1, NOTCH2, FAT1* and several other genes besides *TP53*^12^. Prior attempts to establish a quantitative method for assessing photodamage and skin cancer risk had limited success^17,18^. A method that enables quantitative evaluation of early photodamage is expected to help optimize personalized sun-protective measures and may also serve as a tool for assessing the need and efficacy of early preventative treatment interventions.

In the current work we developed an ultra-deep sequencing-based method to identify CMs in clinically normal epidermis and show differences in CMs between sun-exposed and non-sun-exposed skin areas. We then correlated CMs with skin cancer burden in another independent cohort of cSCC patients and found mutational features in normal skin are significantly associated with cancer risk burden.

## Methods

### Samples

A total of 464 normal human skin samples were collected from 13 Caucasian post-mortem donors over the age of 55 years using Roswell Park’s Rapid Tissue Acquisition Program under a Roswell Park approved IRB protocol within 24 hours of death from frequently sun-exposed (SE) sites (left dorsal forearm) and non-sun-exposed (NE) sites (left medial buttock). Exclusion criteria included any visible skin abnormalities in the tissue areas. Eligible donors were identified and clinically normal appearing skin was harvested. Skin samples were kept in tissue preservation medium, Belzer UW cold storage solution (Bridge to Life, USA) at 4°C until processed. All samples that could be processed within 36 hours or less after death were included in the study. The mean age of the donors was 72.3 years (SD: ±8.2 years; range 60-80 years). The male to female gender ratio was 7:6, and 12/13 donors had no history of skin cancer.

The adipose tissue was removed from each human skin sample using sterile scissors. The samples were cut into strips wide enough to harvest 6 mm punches. The epidermis was separated from the dermis by placing the strips in tubes containing 10 ml of 5U/ml Dispase II (Stem Cell Technologies, USA) and incubated at 4°C overnight and at 37°C for 2-3 hours. After Dispase digestion the specimens were placed in a petri dish containing a small amount of 1x DPBS (Corning, USA) and using sterile tweezers, the epidermis was carefully removed from the dermis. Using disposable biopsy punches, 1, 2, 3, 4 and 6 mm diameter epidermal pieces were taken from the epidermal sheets and punched epidermal pieces were placed into a sterile 1.5 mL vials. In addition to the epidermal punches, large bulk pieces of dermis were also removed from the skin samples using a disposable #15 blade and placed into a sterile 1.5 mL vial for use as a germline control.

For the extended cohort of the study, 20 human skin samples were obtained in a de-identified manner from 8 undergoing surgery for cSCC. The mean age of the donors was 77.9 years (SD: ±12.3 years; range 54-92 years). The male to female gender ratio was 1:1. The study was granted exemption by the Yale University Human Investigation Committee (Protocol 1509016421). All individuals had biopsy-confirmed cSCC that was completely excised by Mohs micrographic surgery with intraoperative histologic verification of clear surgical margins. Immediately following excision of cSCC, adjacent normal skin was excised to facilitate surgical repair and samples for sequencing were immediately harvested. From each individual, two skin samples at a fixed linear distance from the cSCC were obtained from the adjacent, sun-exposed, normal skin. One sample was obtained at a distance of 1mm from the cSCC surgical margin, and one at a distance of 6mm from the surgical margin. From four patients, a tumor sample from grossly visible cSCC was also obtained at the time of surgery. All samples were obtained with a 2mm punch biopsy to a depth of approximately 1mm, including epidermis and superficial dermis.

### DNA isolation

DNA samples from the primary cohort were extracted using Purelink^™^ Genomic DNA mini kit (Invitrogen, USA). Epidermal samples were digested using Proteinase K at 55°C heating block overnight following the manufacturers recommendations. For the extended cohort of samples, skin biopsies were similarly digested using Proteinase K and DNA was purified with phenol-chloroform extraction and ethanol precipitation. DNA was eluted with 28 μL of Molecular Biology Grade Water (Corning, USA) for 1 and 2 mm punches or 36 μL of Molecular Biology Grade Water for 3, 4, and 6 mm punches. The isolated genomic DNA was stored at -20°C and the DNA concentration of each extraction was measured using a Qubit fluorometer or Quanti-iT PicoGreen kit (Invitrogen, USA).

### Ultra-deep Targeted Sequencing

The sequencing libraries were generated using the TruSeq Custom Amplicon kit (Illumina, USA) using 10-50 ng of gDNA. Amplicons of ∼150bp (primary cohort) or ∼250bp (extended cohort) in length were designed using Illumina Design Studio Software. Custom oligo capture probes that flank the regions of interest were hybridized to the gDNA. A combined extension/ligation reaction completed the region of interest between these flanking custom oligo probes. PCR was then performed to add indices and sequencing adapters. The amplified final libraries were cleaned up using AmpureXP beads (Beckman Coulter). Purified libraries were run on a Tapestation DNA1000 screentape chip to verify desired size distribution, quantified by KAPA qPCR (KAPA Biosystems) and pooled equal molar in a final concentration of 2 nM. Pooled libraries were loaded on an Illumina HiSeq Rapid Mode V2 flow cell following standard protocols for 2×100 cycle sequencing (primary cohort), or Illumina NextSeq for 2×150 cycle sequencing (extended cohort).

### Bioinformatics analysis

High quality paired-end reads passing Illumina RTA filter were initially processed against the NCBI human reference genome (GRCh37) using public available bioinformatics tools ^19,20^, and Picard (http://picard.sourceforge.net/). The coverage quality control required at least 80% of the targeted region covered by a minimum of 1,000X coverage. Putative mutations, including single nucleotide variants (SNVs) and small insertions/deletions (Indels), were initially identified by running variation detection module of Strelka^21^ on each SE or NE epidermis sample paired with the matched dermal sample. From the detected SNVs, dinucleotide variants (DNV) or cluster of single nucleotide variants (CSNV) were recognized by running Multi-Nucleotide Variant Annotation Corrector (MAC) ^22^ on the original sequences. The putative mutations detected from all samples were consolidated into a list of unique mutations. Every unique mutation was re-visited in all samples to calculate the numbers of mutant/wildtype reads, as well as variant allele frequency (VAF) in each sample as previously described ^13^.

To distinguish mutations from background errors, we modelled each mutation’s background error rate distributions using VAFs from all control (dermal) samples. For each mutation, we started by fitting a *Weibull* distribution to VAFs from all control samples following a previously published method^23^, then every SE or NE epidermal sample’s VAF was compared to the fitted distribution. A positive sample was defined as the sample’s VAF of a mutation was significantly above background (p < 0.05, after Bonferroni correction). In the extended cohort where the control samples were not available, we adapted a dynamic control strategy, based on the assumption that any somatic mutation cannot be recurrent in more than 10% of all samples at the same site. In the previous primary cohort, all recurrent mutations were within 5% of all samples. For each potential mutation, we first cluster the VAFs of the mutation in all samples. Subsequently started from the cluster with lowest VAF, we transferred all samples of each cluster to the control cohort until at least 90% of all samples are in the control cohort. After mutation calling, all identified mutations including SNVs, DNVs, CSNVs and Indels were annotated using a customized program with NCBI RefSeq database.

Cumulative Relative Clonal Area (CRCA), defined as the overall percentage of biopsied skin area covered by UV-signature mutations (USMs) in a patient skin punch, was calculated as following:

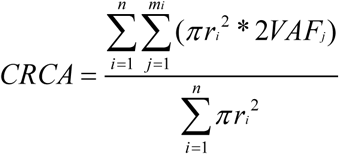

with n = the total number of punches collected in the patient; r_i_ = the size (radius) of each punch; m_i_ = the number of mutations in punch i; VAF_j_ = the variant allele fraction of a specific mutation j. Here, the calculation of CRCA was based on the assumption that all mutations occur in one chromosome of regular diploid genomic regions. Additionally, although we did not consider the situation when multiple mutations occur in the same cell, we did identify mutations that occur on the same reads and combined them into one mutation using MAC ^22^.

### Statistics

The overall mutation numbers and VAFs between two groups, including SE and NE in the primary cohort, and the high- and low-cSCC burden in the extended cohort, were evaluated using a Wilcoxon test. Group-specific markers, including mutations, genes, regions and signatures were identified using a Fisher’s exact test where the two variables in the contingency table were the samples’ sun-exposure status (SE vs NE, in cohort #1) or cSCC burden (high vs low, in cohort #2) and mutational status. Multiple testing correction was implemented using the FDR approach as indicated.

## Results

### Ultra-deep sequencing of epidermal samples using customized focused panels

To generate a focused sequencing panel targeting the most commonly mutated sequences in normal human skin, we selected an area of focus based on a previous dataset^12^. All previous mutations were assigned to 100-bp genomic segments. After sorting the segments by number of mutations, we designed a panel to capture the top 55 most frequently mutated segments from 12 genes (5.5 kb in total, **Table S1**). The majority (65%) of the targeted segments came from the following 3 genes: *NOTCH1, NOTCH2*, and *TP53*. When summarized by coding regions, 79% of the targeted segments lie in protein-coding regions, and the remaining segments were mostly in introns. In the previous dataset^12^, 87% of the samples harbored at least one mutation within this panel. Thus, as designed, this panel captured the most frequently mutated genomic regions in sun-exposed skin, and was highly focused for efficient deep-sequencing to identify low-frequency mutations.

The primary cohort was sequenced using the focused panel in two batches. We first sequenced a discovery cohort of 374 human skin samples from 13 post-mortem donors: 360 epidermal samples, equally acquired from both sun-exposed (SE) and non-sun-exposed regions (NE) using 1 mm, 2 mm, 3 mm, 4 mm or 6 mm punch sizes. From the same 13 donors, DNA from bulk NE dermis (n=14, 1 donor contributed 2 samples) was isolated for germline controls. After initial analysis to determine the optimal punch size, we then tested a separate validation cohort of 90 epidermal samples from 9 of the 13 donors using the most effective punch size (2 mm, as detailed in results “Optimization of punch size for USM detection”). In total, the dataset contains 464 samples: 225 SE, 225 NE, and 14 dermal samples as controls **(Table 1)** from 13 individuals. After sequencing, 85% of samples reached a minimum of 10,000X coverage in at least 80% of the targeted region. The median of average coverage across all samples was 64,730X **(Table S2a)**, with only one sample exclusion (NE sample) due to sequencing failure. This unique design of ultra-deep sequencing from individual matched SE/NE samples enabled us to discriminate between the mutational profiles of SE and NE skin samples.

**Table 1.**
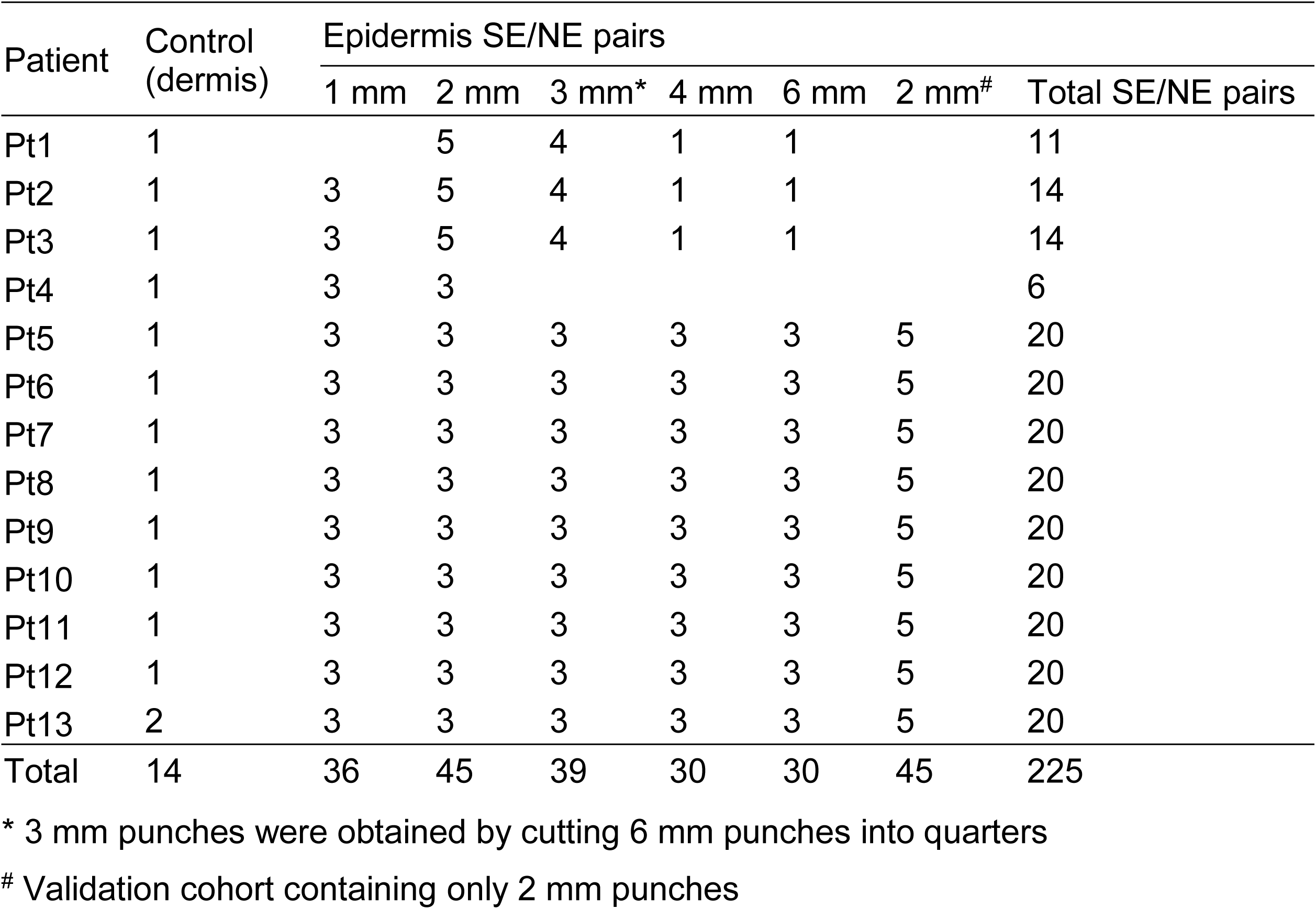
Patient and sample cohort

To better define the clinical relevance of CMs, we sequenced an extended cohort of sun-exposed skin samples from human patients with cSCC. Twenty 2mm punch biopsy specimens were obtained from surgically excised skin from 8 individuals, including 16 normal skin samples and 4 samples of cSCC. For this extended cohort, a custom sequencing panel was designed to encompass the complete protein-coding region of 12 genes with frequently reported mutations in UV-exposed skin (*NOTCH1, NOTCH2, NOTCH3, TP53, CDKN2A, BRAF, HRAS, KRAS, NRAS, KNSTRN, FAT1*, and *FGFR3*), and 1 control gene without expected functional significance in skin (*VHL*). This sequencing panel encompassed 59.5 kb. After sequencing, all samples have at least 80% of the targeted region covered by a minimum of 10,000X coverage. The median value of average coverages across all samples was 47,158X **(Table S2b)**. This extended cohort from cSCC patients would allow us to correlate the features of CMs to patient clinical outcomes.

### Delineate the mutational patterns associated with UV exposure

To identify the mutations solely caused by UV exposure, we characterized the mutational profiles of individual-matched SE/NE epidermal samples. Additionally, we compared the epidermal samples to patient-matched dermal samples followed by an in silico error suppression to remove germline polymorphisms and low-frequency technical artifacts. Dinucleotide and other complex mutations were identified by re-visiting the raw reads using a program that we previously developed ^22^. Altogether, a total of 638 mutations were identified, predominantly single nucleotide variants (SNVs, n = 614 or 96.2%) or dinucleotide variants (DNVs, n = 20 or 3.1%) **(Table S3)**. The median variant allele frequency (VAF) of all mutations was 2.1% (range 0.1% - 36.6%), and only 3% mutations reached a VAF greater than 10%.

Among the 55 targeted genomic segments, mutations were detected in 50 segments with an average of 7.1 and 4.7 mutations per segment in SE and NE samples, respectively **(Figure 1a)**. Two segments were significantly (FDR p<0.001) associated with UV-exposure status, approximately corresponding to *TP53* amino acids 227-261 (“*TP53-3*”, mutations in SE/NE = 38/0) and *NOTCH1* p.449-481 (“*NOTCH1-9*”, mutations in SE/NE = 30/4). Interestingly, mutations in an adjacent region in *NOTCH1* p.419-449 (“*NOTCH1-10*”) were not associated with UV exposure (mutations in SE/NE=48/40), even though “*NOTCH1-10*” was the most frequently mutated segment in the current study. Additionally, mutations were marginally enriched in SE samples (FDR p<0.1) in three other segments: two in *NOTCH1* (“*NOTCH1-14*” and “*NOTCH1-19*”) and one in *GRM3* (“*GRM3-2*”). On the gene level, mutations in SE samples were only significantly enriched in *TP53* (FDR p<0.001), and marginally significant in *GRM3* (FDR p<0.1). Overall, the numbers of mutations in SE samples were 6.3 times higher than NE samples in *TP53*, and 4.3 times in *GRM3* **(Figure 1b)**. Mutations identified in nine other genes did not exhibit significant association with sun-exposure status either on the gene- or segment-level: *NOTCH2, ARID1A, SALL1, SCN1A, ERBB4, FAT4, FGFR3, ADGRB3 and PPP1R3A*. These findings indicate a highly genomic-region-specific pattern of the accumulation of UV-induced somatic mutations.

**Figure 1.**
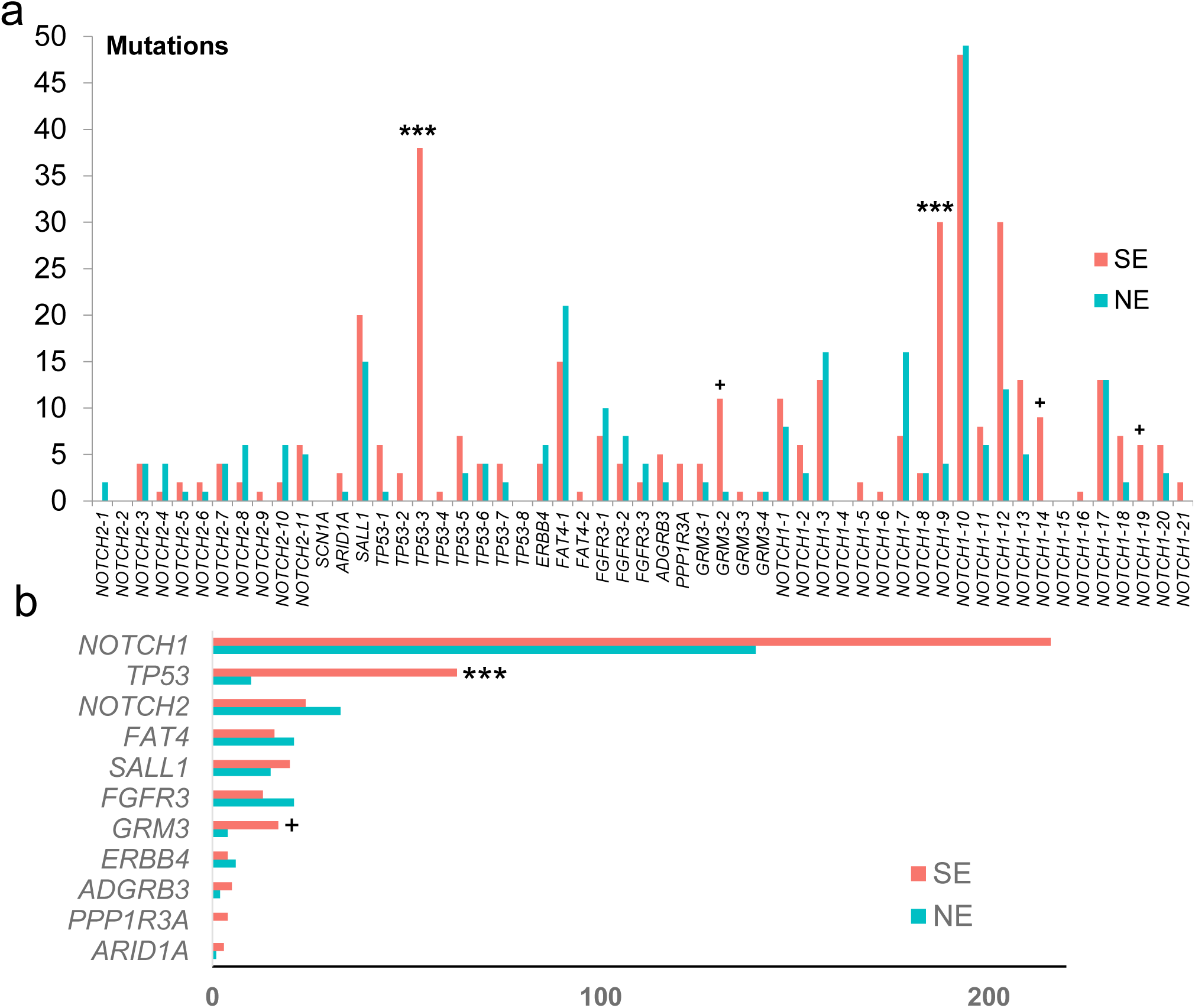
Region-specific enrichment of somatic mutations in sun-exposed skin. a). Graph shows the number of mutations identified within each 100-bp genomic target window grouped by SE and NE skin types. b). The overall gene-level number of mutations from SE and NE samples. Stars indicate the segments or genes where mutations are significantly enriched in the SE samples (FDR p values: *** p<0.001; ^+^ p<0.1).

We next investigated potential hotspots and mutations associated with UV-exposure. After sorting all mutations by their genomic locations, one specific region in *TP53* (p.217-280), appeared to be “mutation exempt” in comparison to surrounding regions in NE samples. In contrast, this region was highly mutated in SE samples **(Figure 2a)**. We reanalyzed a recent study involving RNASeq of both SE and NE normal skin samples^11^, and found four mutations in this region, all from SE samples **(Table S4)**. To identify mutations associated with UV exposure, we focused on highly recurrent mutations (present in ≥ 5 samples, n = 18). By comparing the frequency in SE and NE skin samples, we identified six mutations significantly enriched in SE samples: *TP53* R248W, *NOTCH1* P460L, *NOTCH1* S385F, *NOTCH1* E424K, *TP53* G245D and *NOTCH1* P460S, and nearly all of them were exclusively found in SE samples (FDR p<0.05, **Figure 2b**). No mutation was significantly enriched in NE samples. Five of the six SE-enriched mutations were found in both discovery and validation cohorts, indicating they were unlikely to be caused by batch-effect. Unexpectedly, one specific mutation (*NOTCH1* E424K) was associated with significantly elevated VAFs (median = 10%, p<0.001, Wilcoxon test), about five-fold higher than other mutations (median VAF = 2.1%, **Figure 2a, 2b**). Through protein structure modelling **(Figure 2c)**, we found that the *NOTCH1* E424K mutation is predicted to disrupt the binding of *NOTCH1* to delta-like canonical ligand 4 (*DLL4*), a negative regulator of the Notch signaling pathway^11^. By prohibiting formation of a salt bridge between *NOTCH1* E424 and *DLL4* K189/R191, the mutation E424K creates a repulsive force that inhibits *DLL4* binding ^24^. Based on the biological role of *DLL4* and *NOTCH1*, the *NOTCH1* E424 mutation is expected promote epithelial proliferation ^25,26^. The overall prevalence of the *NOTCH1* E424K mutation in our dataset is 2.7%. For comparison, in GENIE cBioPortal^27^, *NOTCH1* E424K is mutated in 1.3% of cutaneous SCCs, 0.04% in melanomas, and is rarer in other cutaneous or non-cutaneous malignancies **(Table S5)**.

**Figure 2.**
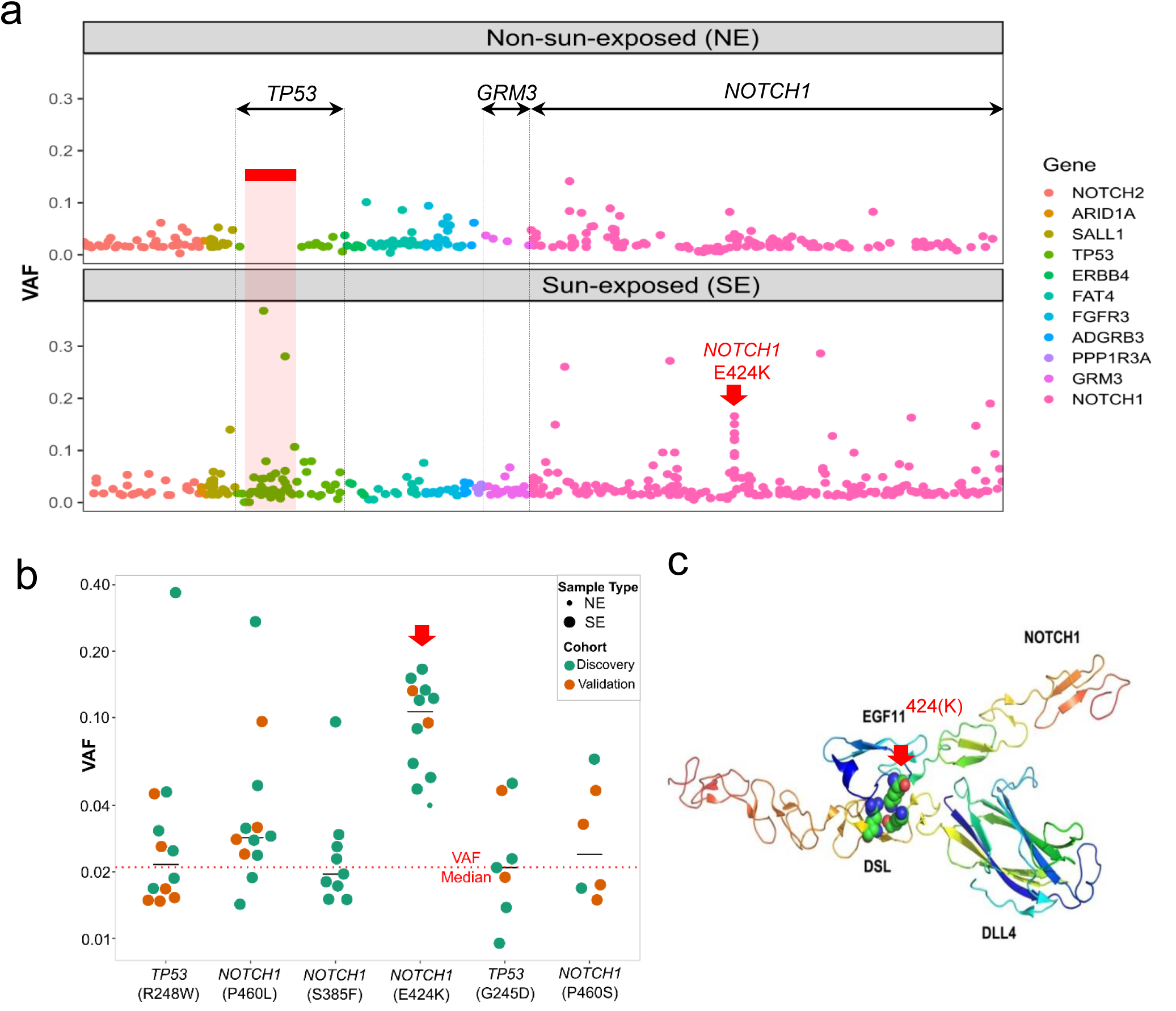
Hotspots and mutations associated with UV-exposure. a). All mutations are ordered by their genomic locations. X-axis: the order of the mutation’s genomic location. Y-axis: variant allele fraction (VAF) of individual mutations. Color depicts the gene harboring the mutations. The three genes demonstrating significant difference between SE and NE, either on the gene level or segment level, were labeled on top (*TP53, GRM3, NOTCH1*). One specific mutation with elevated VAFs (*NOTCH1* E424K) is indicated with a red arrow. b). The VAF of the six individual mutations that are significantly enriched in SE vs NE epidermis in the primary discovery (green) and validation (brown) data sets. The dotted red line represents median VAF of all mutations and black lines indicate the median of each group. c). The predicted protein complex structure of NOTCH1 and DLL4 to show the position of the mutant E424K and the interacting partners, DLL4 K189/R191, in wild type.

### UV-signature mutations exclusively account for the elevated mutation burdens in SE skin

We next intercorrelated the identified mutations with previously known UV-signature mutations (USMs), i.e., C>T transition at dipyrimidines ^4^. Among all 638 mutations in SE and NE samples, 298 were USMs. Of these 298 USMs, 76% were present in SE samples. USMs were significantly enriched in SE compared to NE samples (n = 226 and 72, respectively, p<0.001, Fisher’s exact test). Especially among the high-VAF mutations, 18 of 19 mutations with VAFs above 0.1 were from SE samples, and most (13 of 18) were USMs. Conversely, non-UV-signature mutations (NUSMs) were present’ approximately equally (n= 159 and 181, ns, Fisher’s exact test) in SE and NE skin types **(Figure 3a)**, indicating that these mutations may not be directly associated with UV-exposure.

**Figure 3.**
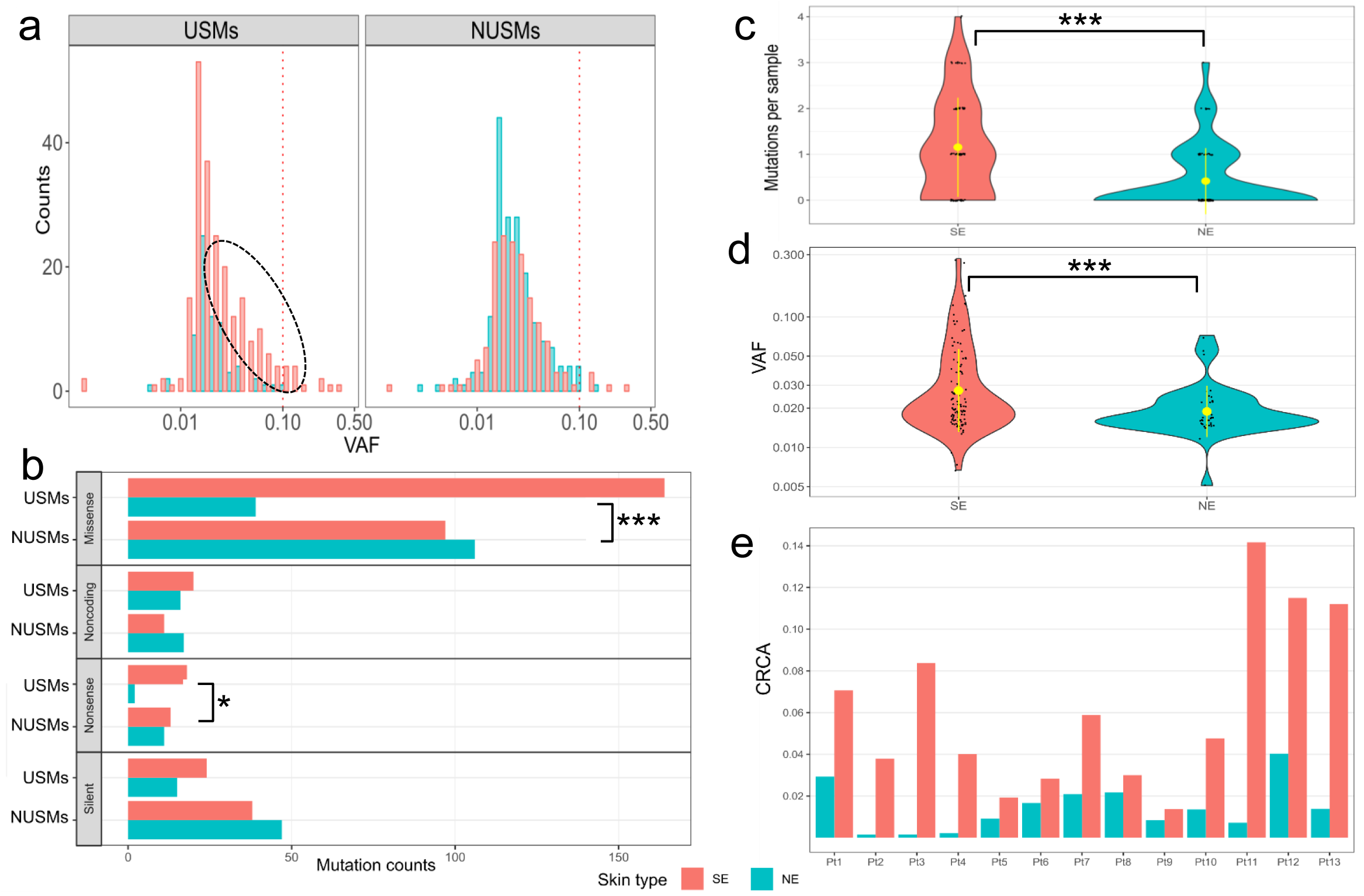
UV-induced DNA damage assessed by USMs. a). Only UV-signature mutations are associated with sun-exposure status. Left: higher numbers of USMs are found in SE than NE skin. Right: NUSMs are almost equally presented in SE and NE samples. Red dotted line indicates high-VAF (>0.1). Black dotted circle indicates extra USMs in SE skin compared with NE skin. b). The numbers of mutations by each amino-acid-change type found in SE and NE skin, grouped by USMs and NUSMs. Overall distribution c.) of the numbers of USMs per sample and d.) the VAFs of the mutations using the 2 mm punch size. Inside the violin plots: black dots – original data points from individual samples; yellow dot with bar - averaged value with standard deviation. SE samples are associated with higher numbers of USMs, as well as higher VAFs indicating potential larger clones. e). Cumulative Relative Clonal Area (CRCA) was developed to represent the overall percentage of the biopsied skin areas that are covered by clonal mutations. In all 13 current individuals, CRCAs were higher in SE than in the matched NE group, with the ratios of SE/NE ranged from 1.4 to 55.0 (mean = 11.2). Statistical tests used: figure 3b, Fisher’s exact test with multiple test correction implemented using the FDR method; figure 3c, 3d: Wilcoxon test; *p<0.05, **p<0.01, ***p<0.001.

To explore specific community enrichment patterns in different mutational function groups, we classified all 638 mutations into four effect-groups: nonsense, missense, silent and noncoding. Inside each effect-group, we correlated the mutational properties (USM vs NUSM) with the matched samples’ sun-exposure statuses (SE vs NE) **(Figure 3b)**. Significant enrichment of USMs were observed in two of four effect-groups by Fisher’s exact test: nonsense (FDR p<0.05) and missense (FDR p<0.001). Specifically, nonsense mutations were 9 times more frequently occurring in SE skins than in NE skins, and similarly enriched by 4.2 times for missense mutations. These findings indicate that the mutations initiated by UV radiation are further selected by the host system or inter-clonal competition ^28^, in which the mutations with functional impacts give the host clone greater fitness.

### Quantification of UV-induced DNA damage level by UV-signature mutations

We next investigated the feasibility of using CMs to quantify UV-induced DNA damage. This was based on the hypothesis that SE samples harbor more CMs and are associated with higher VAFs compared to NE samples. Since our analyses indicated NUSMs were not correlated with UV exposure, only USMs were used for quantifying UV-induced DNA damage. To avoid the potential bias introduced by different punch sizes, initially only the most abundant size of 2 mm (n = 90 and 89, SE and NE, respectively) **(Figure 3c)** was analyzed. A three-fold difference was observed in the average USMs per sample between SE (mean = 1.2) and NE (mean = 0.4), which was significantly higher (p <0.001, Wilcoxon test). Multiple USMs were found in 33% of SE samples but only 9% of NE samples **(Table S6)**. Additionally, the identified USMs had significantly higher VAFs in SE (mean = 3.7%) than NE (mean = 2.1%) samples, (p < 0.001, Wilcoxon test), indicating the presence of larger clones in SE samples **(Figure 3d)**. We further extended the analysis to include all punch sizes, and found the pattern was consistent with 34% of SE and only 6% of NE samples having multiple USMs and three-fold higher average USMs per sample in SE (1.0) than NE (0.3) samples (p < 0.001, Wilcoxon test). These findings of increased USMs and elevated VAFs in SE than NE skin would then serve as the cornerstones for the quantification of UV-induced DNA damage.

In order to overcome the heterogeneity between samples, we developed Cumulative Relative Clonal Area (CRCA) as a single metric to assess the overall patient-level burden of CMs. The CRCA was defined as the overall percentage of biopsied skin area covered by USMs in a patient skin punch, which account for both the number of USMs and their VAFs **(Figure 3e)**. It is worth mentioning that our data did not allow us to distinguish whether mutations occurred independently or were present in the same clone. Hence, CRCA does not provide an exact measure of the mutated cell population, but rather serves as an index of the mutation burden in the sampled area. To minimize the potential chance for repeated counting of co-occurring mutations in the same cells, co-occurring mutations were identified, primarily dinucleotide CC>TT mutations, and consolidated. When counted separately by sun-exposure status, the median CRCA across the 13 patients was 6.1% (range 1.4-14.2%) in SE and 1.4% (range 0.1-4.0%) in NE sites. On individual patient level, the CRCAs were higher in SE than the matched NE skin in all patients, with the average ratio of 11.2-fold higher (range = 1.4 - 55.0-fold). These CRCAs were calculated using only USMs. If all CMs were included, the CRCA would be only 2.2-fold higher (range = 0.8 - 5.6-fold) in SE than NE skin (data not shown). Based on these results, CRCA may have the potential to be used as an objective measurement of the level of UV-induced DNA damage.

### The effect of punch size on USM detection

In the discovery cohort, we sought to evaluate different punch sizes to determine the most efficient one for detecting USMs. Theoretically, although larger punches likely contain more clones, they tend to become less effective for detecting smaller clones due to a dilutional effect by other clones harboring no or different mutations **(Figure 4a)**. Overall across all five punch sizes, USMs were detected in 54% of the SE, which was significantly higher than the 21% of the NE (p < 0.001, Fisher’s exact test). Between different punch sizes, 2 mm punches were found to have the highest positive rate of 64%, and with the most significant difference between SE and NE samples (p < 0.0001, **Figure 4b**). Thus, only 2mm punches were collected in the 90-sample validation cohort and the extended cohort from cSCC patients. In the validation cohort, similarly, we found the SE samples had higher numbers of USMs and the positive rate of USMs (69%) was similar to the discovery cohort (64%).

**Figure 4.**
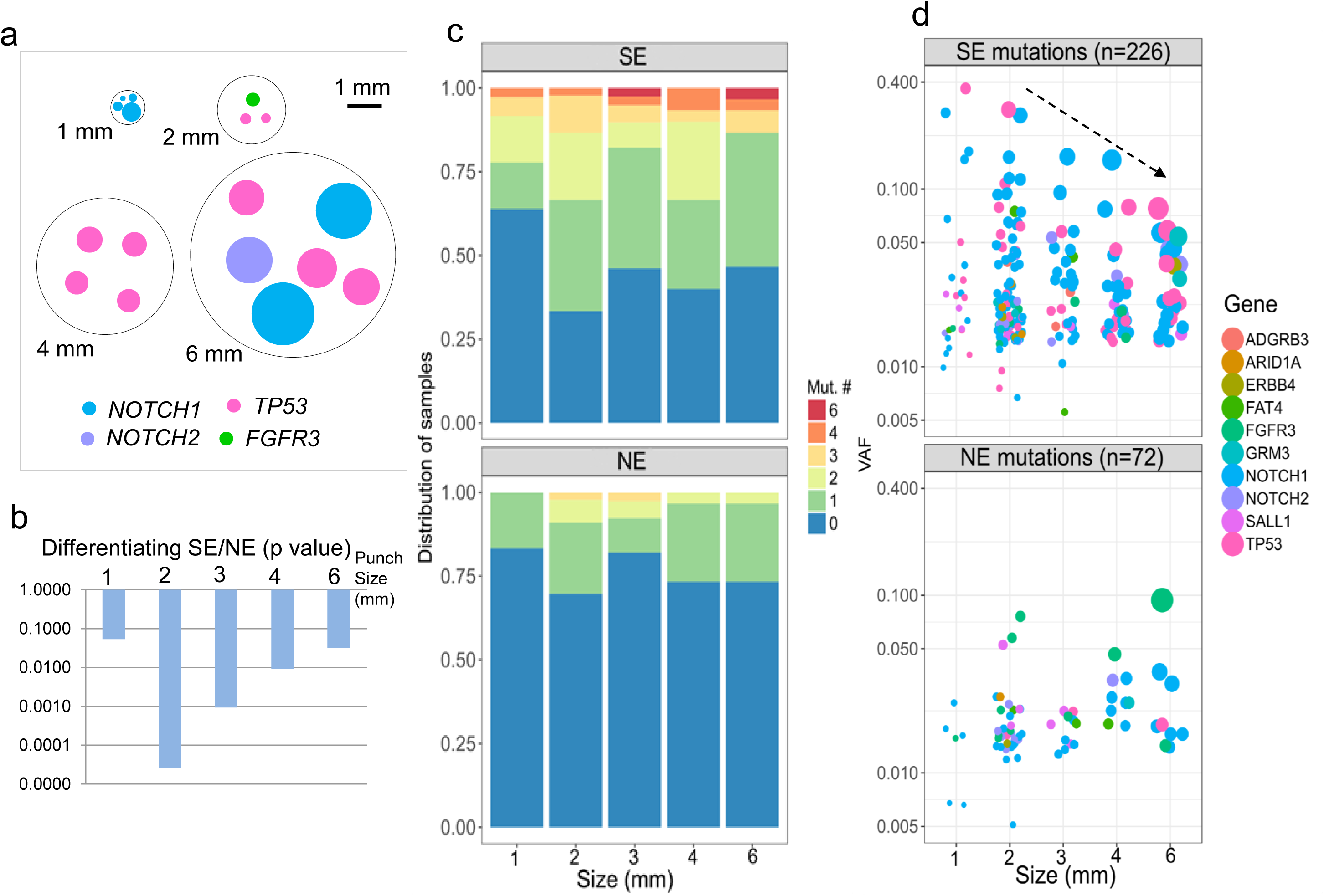
Optimization of punch size for detecting USMs. a). A representative figure showing one representative punch of each collection size. We selected the sample with the highest number of mutations under each size for easy illustration. Every mutation is plotted as a dot with its size calculated to match the clonal area harboring the mutation. One punch size, 3 mm was not shown as it was obtained by cutting a 6 mm punch into quarters. b). In the discovery cohort, 2 mm was found to be the most efficient size in differentiating CRCA from SE and NE skin samples by p value. c). Distribution of numbers of USMs per sample at each punch size, after combining both the discovery and validation cohorts. d). VAF of USM detected in different size punches. The size of the dot indicates the approximate relative area of cells containing the mutation. In SE samples, VAFs of USMs detected from larger punches are associated with smaller variations.

When combining the discovery and validation cohorts, the SE samples had the highest positive rate of 67% for USMs in 2 mm samples and were significantly higher than NE samples (p < 0.001), followed by 60% in 4 mm (p < 0.05), and 54% in 3 mm (p < 0.05). Interestingly, the USM positive rates were relatively lower in the largest punch size of 6 mm (53%) and the smallest 1 mm (36%). In all NE samples, positive USM rates ranged from 17-30% (**Figure 4c**). Moreover, the punch size also affected the detected VAFs of the mutations. Specifically, in SE samples, larger punches were associated with smaller VAFs. The VAFs’ standard deviation was the highest in 1 mm punches (8.9%) and decreased with punch size: 2 mm (4.3%), 3 mm (2.8%), 4 mm (2.6%) and 6 mm (1.7%). This trend, between VAF range and punch size, was not present in NE samples **(Figure 4d)**. These results suggested that the most effective punch size in detecting USMs under the current sequencing condition was 2 mm.

### Mutation nucleotide contexts enriched with UV-exposure

We next assessed the enrichment of different mutation nucleotide contexts in SE skin. The mutation nucleotide contexts were defined by each SNV’s trinucleotide and DNV’s dinucleotide contexts. A total of 83 contexts were identified from current mutations, including 13 contexts matching to previously described USMs^4^. None of the remaining 70 non-USM contexts were enriched in SE or NE samples **(Figure 5a)**. The 13 previously defined USM contexts were not equally enriched in SE samples. After multiple test correction, only 5 of the 13 contexts were significantly enriched in SE samples (FDR p<0.05), including the dinucleotide CC>TT context, which was exclusively found in SE samples **(Figure 5b)**. The most significant mutation context enriched in SE samples was T[C>T]C (FDR p = 0.00013), which was in consonance with the previously defined “Mutational Signature #7” in skin cancers ^29^. The remaining eight UV-signature contexts were not significantly enriched in SE samples. Of particular note, G[C>T]C, which was the most abundant context by total number of mutations, appeared to be equally presented in SE and NE skin samples and therefore not associated with sun-exposure.

**Figure 5.**
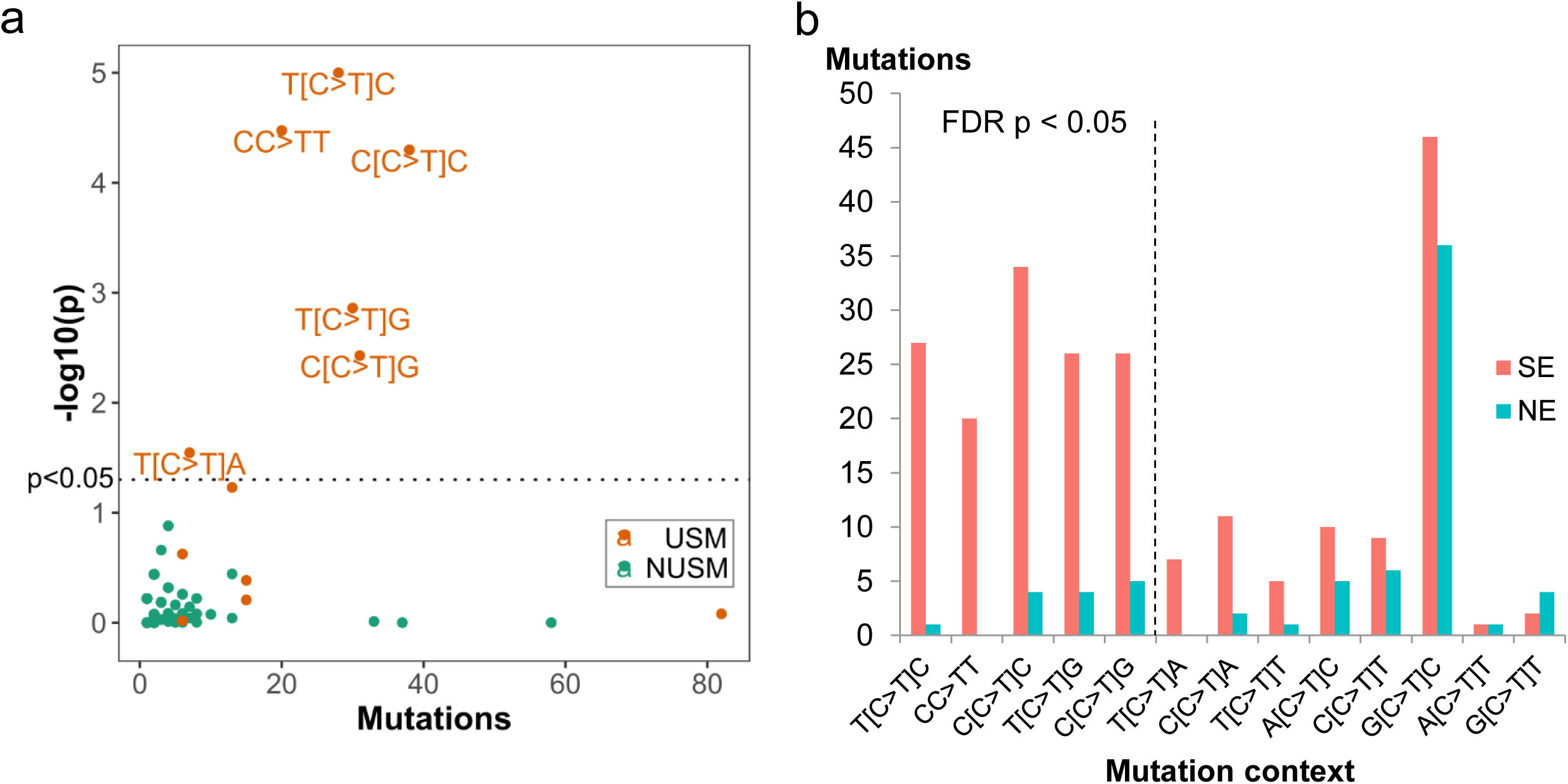
Mutational contexts associated with UV-exposure. a). Each dot represents a specific mutation context of SNVs and DNVs. X-axis: the total numbers of mutations of each context; y-axis: p value of the context for differentiating SE and NE skin, shown as -log(p). The dotted line indicate p<0.05 (the above area). None of the NUSM contexts was significant. b). Further refinement of USM contexts by depicting the numbers of mutations in SE and NE skin for all current USM contexts. Mutation contexts are ordered by the p value of SE vs NE in an increasing order from left to right. Multiple test correction was implemented using the FDR method. The dotted line indicates FDR p<0.05 (the left side).

### Clonal mutations are correlated with cSCC burden

To define the clinical significance of CMs and investigate the potential association with skin cancer risk, we sequenced an extended cohort of 20 samples (16 SE normal skin and 4 cSCC) from eight patients with cSCC using a 59.5 kb customized panel as described above. Four individuals (including 8 normal skin samples and 2 cSCC samples from face, scalp, and arm) had a low burden of skin cancer with only a single diagnosis of cSCC and few AKs (low-cSCC). Four individuals (including 8 normal skin samples and 2 cSCC samples from face, hand, and lower leg) had a high burden of skin cancer with severe UV damage, multiple prior cSCC (range 3-10) and many AKs (high-cSCC). Low-cSCC and high-cSCC patients were matched for age (mean age 76.5 and 79.3, respectively). Normal skin samples were all sun-exposed, and were obtained a linear distance of either 1mm or 6mm from the clear surgical margin of the excised cSCC, allowing for analysis of CMs arising in skin subjected to carcinogenic UV radiation. Visible AKs were not present in normal skin samples. A total of 535 somatic mutations were identified **(Table S7)**, with a median VAF of 1.2%. Only 15 mutations had VAF greater than 10%, most of which (10 of 15) were from the cSCC tumor samples **(Figure 6a)**. The median numbers of mutations per sample in each group were 22 and 17.5 for the high- and low-cSCC normal skin samples (marginally significant, p=0.078, Wilcoxon), and 41.5 for the cSCC samples. The overall mutation rates in normal skin were 0.45 and 0.29 mutations per MB, in high- and low-cSCC patients, respectively. The latter was comparable to the rate of SE normal skin of non-cancer patients in the primary cohort (0.31 mutations per MB), despite the technical differences between the two cohorts such as sequencing depth, targeted regions and punch sizes.

**Figure 6.**
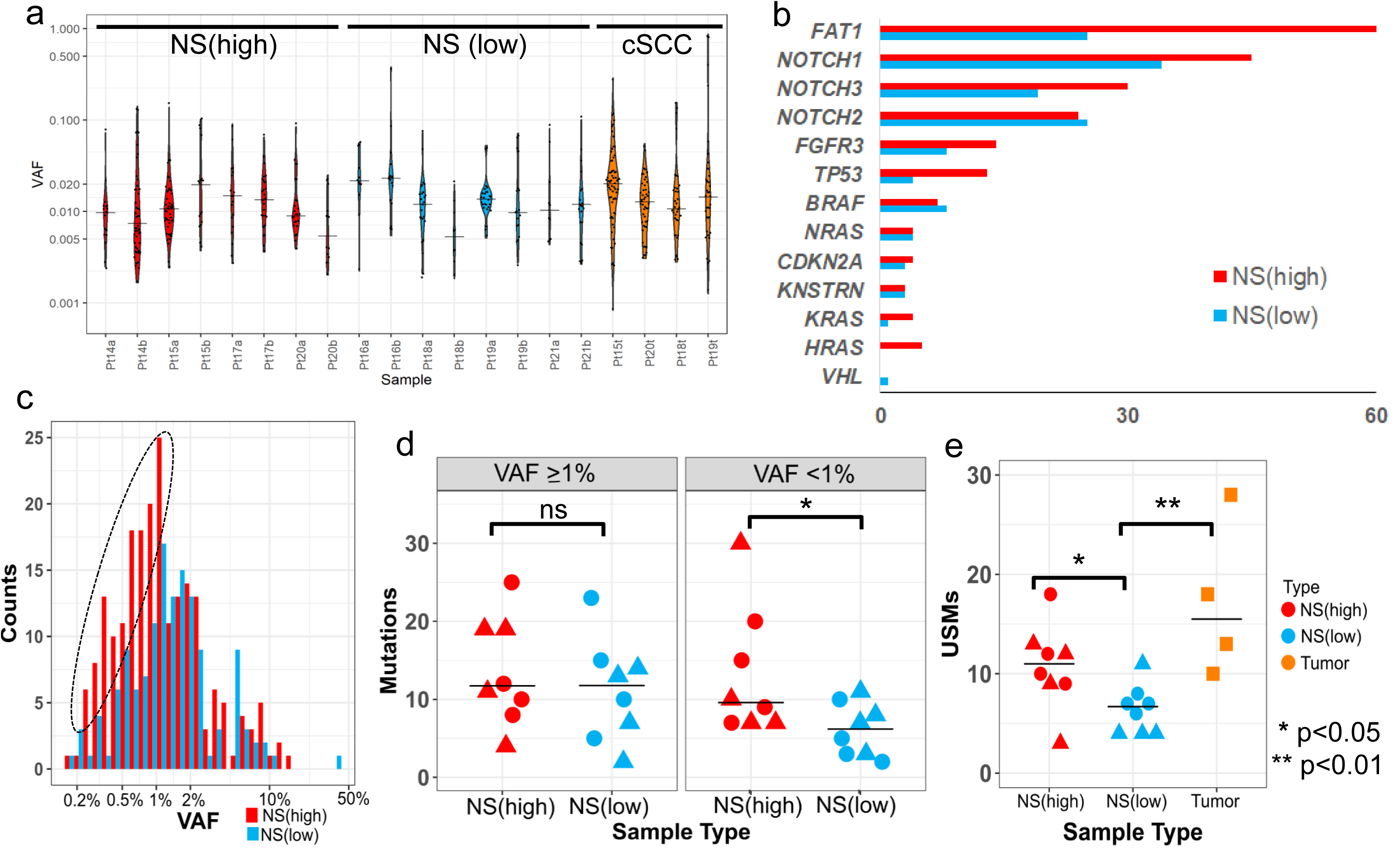
Clonal mutations are correlated with cSCC burden. a). Violin plots depicting the overall distribution of somatic mutations in each sample, ordered by sample type. b). Mutation numbers by genes in the normal skin. NS (high) - normal skin from high-cSCC patients; NS(low) - normal skin from low-cSCC patients. c). High-cSCC patients are associated with increased low-VAF (<1%) mutations. Histogram depicting the distribution of VAFs of the detected mutations in normal skin separated by cSCC burden. The dotted oval highlights the increased low-VAF mutations in the normal skin of high-cSCC patients compared with low-cSCC patients. d) Number of mutations per sample in normal skin, separated by high-(≥1%) and low-(<1%) VAFs; e) Number of USMs per sample in high- and low-cSCC normal skin (NS), and cSCC tumors. Shape indicates the two normal skin samples from each patient, taken either 1mm (circle) or 6mm (triangle) from the surgical margin.

The frequently mutated genes in normal skin (more than two mutations per gene on average) included *FAT1, NOTCH1, NOTCH2, NOTCH3, FGFR3* and *TP53* **(Figure 6b)**. Two of the genes were mutated at least twice as frequently in the normal skin of high-cSCC patients as that of low-cSCC patients: *TP53* (ratio = 3.25) and *FAT1* (ratio = 2.4). Additionally, two less frequently mutated genes, *KRAS* and *HRAS*, were almost exclusively mutated in high-cSCC patients (9 of 10). None of these differences reached statistical significance after multiple test correction (data not shown), indicating that larger cohorts will be needed to further explore these potential associations.

Although the normal skin of high-cSCC patients contain more mutations per sample, unexpectedly, these mutations were associated with significantly lower VAFs (median=1.0%) than the normal skin of low-cSCC patients (median = 1.3%, p = 0.011, Wilcoxon). We found this overall reduction in VAF resulted from a higher number of low-frequency mutations in high-cSCC patients **(Figure 6c)**. For mutations with VAF greater than 1%, the mutations were equally present in high- and low-cSCC patients. However, for low-VAF mutations (defined as <1%), the numbers of mutations per sample were significantly higher in high-cSCC (median = 9.5) than low-cSCC patients (median = 6, p = 0.032, Wilcoxon, **Figure 6d**).

We next further refined the analysis by focusing on USMs. There were a total of 206 USMs, including 8 CC>TT DNVs. We observed a significantly greater number of USMs in the high-cSCC normal skin samples (median = 11) than the low-cSCC ones (median = 6.5, FDR p = 0.015) **(Figure 6e)**. The tumor samples were found to harbor even higher numbers of USMs (median = 15.5). The CRCA values, as defined in the primary cohort, were significantly higher in the tumor than the normal skin samples (FDR p = 0.03) in the extended cohort. The normal skin samples from high-cSCC patients had slightly higher CRCAs (median = 0.37) than low-cSCC patients (median = 0.31), but the difference was not statistically significant (p = 0.16). The CRCA is essentially the sum of VAF values for all detected mutations, normalized for biopsy size. The lack of a significant difference between CRCA values for high-cSCC and low-cSCC skin samples is likely due to the observation that the increased mutations present in high-cSCC samples were enriched for low-frequency mutations (VAF < 1%). We found no significant difference in overall mutation burden, VAF, USMs, or CRCA between normal skin samples collected at 1mm versus 6mm from the surgical margin. Lastly, almost all mutations (>99%) were present only in one of two skin samples from the same patient. The absence of shared recurrent mutations across different samples from the same individual indicates that the identified mutations arose independently.

## Discussion

Most cancers are initiated by accumulation of somatic mutations^30,31^. However, early mutations in normal tissues are difficult to detect due to the low abundance and random patterns. Several recent studies demonstrated the feasibility of detecting clonal mutations (CMs) using high-throughput sequencing in various tissue types^11,12,32^. However, the contribution of these CMs to cancer remains unclear in several ways: how they are generated, what types of mutations are generated by which exogenous and endogenous carcinogens; how the CMs are accumulated and selected by the host microenvironment and inter-clonal competition ^28^; and which mutations contribute or lead to the development of cancer. Indeed, all types of tissues are under the influence of multiple intrinsic and extrinsic factors that vary greatly by individual’s lifestyle and environment. Therefore, studying the CMs generated by one specific carcinogen requires comparative studies of matched sample types.

To the best of our knowledge, the current study of paired SE and NE skin areas is the first analysis of individual-matched normal human skin to specifically characterize UV radiation’s mutational effects. We optimized our detection of UV-induced CMs by: 1) acquiring matched SE/NE skin samples from the same individual to control for aging and other environmental factors unrelated to UV; 2) separating epidermal from dermal layers to decrease non-epidermal background DNA quantity; and 3) ultra-deep DNA sequencing for maximized sensitivity followed by error-suppression to exclude sequencing and alignment errors. Consistent with previous studies ^11,12^, CMs were widespread in epidermal samples. As expected, mutation burden and VAFs were significantly elevated in SE samples. The mutational signatures of the current CMs are consistent with those previously found in skin cancers ^29^, supporting the contribution of the CMs to potential ongoing tumorigenesis. Markedly, our unique approach allowed us to gain several important new insights about epidermal CMs. First, we identified the existence of “mutation-exempt” regions in human genomes. Although mutations frequently occur across most of the sequenced regions in NE skin, presumably due to metabolism and aging related factors, no detectable mutations were found in these mutation-exempt regions. It is unclear whether the absence of mutations in these genomic regions is caused by an active protection or a passive selection mechanism involving altered clone fitness. Interestingly, the “mutation-exempt” property of these regions appears to be altered upon exposure to UV radiation, and these regions become highly mutable. Further studies are warranted to explore how this mechanism is abrogated by UV radiation. Second, USMs were significantly enriched in Glutamate Metabotropic Receptor 3 (*GRM3*) in SE skin, which was previously identified as a potential therapeutic target in melanoma ^33^, but not reported as a cancer driver in cutaneous SCC. Third, we identified six mutations that were almost exclusively mutated in SE skin. All six mutations had been previously reported in human cutaneous squamous cell carcinomas in the cBioPortal ^27^. Among these mutations, *TP53* R248W and G245D were highly recurrent with hundreds of occurrences reported in *COSMIC* ^34^, indicating that the presence of these mutations may be representative of an early phase of carcinogenesis.

Consistent with the current finding that UV-exposure results in higher USM burden, and the known knowledge that UV-exposure directly correlates with the risk of cSCC ^35^, the results of our extended cohort of cSCC patients provided direct evidence that elevated USM burdens are associated with increased burden of cSCC. Presumably, this burden correlates with risk of future cSCC as well. Unexpectedly, we further discovered that most mutational difference between normal skin of high- and low-cSCC patients derived from low-frequency clones (VAF<1%) but not the “expanded” clones (VAF≥1%). It remains unclear why such difference was not seen in the expanded clones. One potential explanation is that the expanded clones might be under more aggressive immune surveillance, as it has been previously reported that the immune system preferably targets larger clones than smaller ones ^36^. The low-frequency clones, on the other hand, are less actively monitored by the immune system and may more truthfully represent the level of ongoing mutational activity or genomic instability. In any case, the total USM burden in sun-exposed skin of patients with cSCC may be a more accurate measure of skin cancer risk than VAF or clonal area.

Our approach was directed by future clinical utilities, focusing on quantitative measurement of UV-induced DNA damage for sun-protection, and cSCC patient risk stratification. These results demonstrate the feasibility of using a small panel of genomic regions (5.5 kb) to quantitatively measure UV-induced CMs. We established Relative Cumulative Clonal Area (CRCA) as a combined measure of mutation burden and relative abundance, which was strongly correlated with sun exposure status, but not with cSCC burden in sun-exposed skin. In the current study, we found the most effective punch size for capturing CMs was 2 mm, which is also clinically favorable as it leaves relatively smaller scars due to the small diameter punch. In future, a non-invasive skin sampling method may provide even wider accessibility to epidermal sampling. In addition, the efficiency of this panel is related to the performance of sequencing method and mutation calling algorithm, which will likely be improved with adoption of more sensitive future methods focusing on the genomic hotspots that are sensitive to UV exposure.

The current study focused on the most frequently mutated regions in sun-exposed skin samples defined by the mutations in a previous study ^12^. However, we note that many of these regions are mutated in both sun-exposed and non-sun-exposed skin samples, indicating that many mutations in these regions were unrelated to UV exposure. In fact, only 6 of 55 original regions were found to harbor significantly enriched mutations in SE samples. Future studies, including much larger targeted regions, are needed to systematically identify UV-sensitive genomic regions. The skin samples were collected at the same time; therefore, they do not provide longitudinal information about clone initiation and progression. While our analyses of the extended cohort indicate that the burdens of CMs in normal skin are correlated with cancer risk in cSCC patients, this finding needs to be validated in a larger cohort of patients. Future studies including biopsies of both SCC and adjacent normal skin acquired at multiple time points are warranted to unveil the complete role of these CMs in cancer.

## Conclusions

In summary, this study revealed previously unknown mutational patterns associated with UV-exposure, providing important insights into the early carcinogenic effects of UV radiation. The quantification of CMs has the potential to become a cornerstone for future development of quantitative measures of UV-induced DNA damage, as measured by CRCA, in the clinical setting to monitor early carcinogenesis and highlight the importance of sun protection. The identified association between cSCC burden and mutation status, especially low-frequency CMs, if validated in an expanded cohort, may become a novel biomarker for risk stratification of cSCCs.

## Ethics Statements

All specimens in the primary cohort were collected from post-mortem donors collected in collaboration with Buffalo’s local organ procurement organization (ConnectLife, formerly Unyts) the Roswell Park’s Rapid Tissue Acquisition Program under a Roswell Park approved IRB protocol. Specimens in the expanded cohort were collected from discarded surgical tissue under a Yale University Human Investigation Committee approved protocol.

## Availability of data and materials

The datasets used and/or analyzed during the current study are available from the corresponding authors upon request.

## Acknowledgments

The authors thank the excellent technical help provided by Paula Pera, MS, assistance with design of the 13-gene custom sequencing panel for the extended cohort provided by Yuemei Zhang, MD (Yale University School of Medicine), and assistance with library generation and sequencing of the extended cohort provided by Mei Zhong, PhD (Yale Stem Cell Center).

We dedicate our work to Dr. Oscar Colegio, who passed away suddenly on June 14th, 2020. Oscar was not just a colleague and co-author, he was a passionate and exceptionally empathetic physician, a brilliant researcher, a thoughtful friend. Oscar was among the leading transplant dermatologists in the world. He had a captivating personality and a unique ability to connect ideas and people. This manuscript is a testament to Oscar’s ability to bring people together. As we will continue our collaboration Dr. Colegio’s insight, mentorship, wit and hard work will be greatly missed.

## List of abbreviations

UV: Ultraviolet
CM: Clonal mutation
NMSC: Nonmelanoma skin cancer
SE: Sun-exposed
NE: Non-sun-exposed
USM: UV-signature mutation
NUSM: Non-UV-signature mutation
CRCA: Cumulative Relative Clonal Area
cSCC: Cutaneous squamous cell carcinoma
AK: Actinic keratosis
SNV: Single nucleotide variant
Indels: Insertions/deletions
DNV: Dinucleotide variant
CSNV: Cluster of single nucleotide variant
MAC: Multi-Nucleotide Variant Annotation Corrector
VAF: Variant allele frequency

